# BNMCMC: a software for inferring and visualizing Bayesian networks using MCMC methods

**DOI:** 10.1101/414953

**Authors:** A. K. M. Azad, Salem A. Alyami, Jonathan M. Keith

## Abstract

**Motivation:** Bayesian networks (BNs) are widely used to model biological networks from experimental data. Many software packages exist to infer BN structures, but the chance of getting trapped in local optima is a common challenge. Some recently developed Markov Chain Monte Carlo (MCMC) samplers called the Neighborhood sampler (NS) and Hit-and-Run (HAR) sampler, have shown great potential to substantially avoid this problem compared to the standard Metropolis-Hastings (MH) sampler.

**Results:** We have developed a software called BNMCMC for inferring and visualizing BNs from given datasets. This software runs NS, HAR and MH samplers using a discrete Bayesian model. The main advantage of BNMCMC is that it exploits adaptive techniques to efficiently explore BN space and evaluate the posterior probability of candidate BNs to facilitate large-scale network inference.

*Availability:* BNMCMC is implemented with C#.NET, ASP.NET, Jquery, Javascript and D3.js. The standalone version (BN visualization missing) available for downloading at https://sourceforge.net/projects/bnmcmc/, where the user-guide and an example file are provided for a simulation. A dedicated BNMCMC web server will be launched soon feature a physics-based BN visualization technique.

*Contact:*

## 1 Introduction

Markov chain Monte Carlo (MCMC) methods are powerful tools for inferring Bayesian network (BN) structures from data (Su and Borsuk, 2016), yet a wide range of software packages for BNs do not adopt them. For example, *BNFinder* (Wilczynski and Dojer, 2009) uses an exact algorithm, *bnlearn* (Scutari, 2010) uses constraint-based, pairwise, score-based and hybrid algorithms, *Bayes Server* (Ser, 2017) uses the PC algorithm, and *GeNie* (GeN, 2017) uses Bayesian Search, PC, essential graph search, greedy thick thinning, tree augmented naive Bayes, augmented naive Bayes, and naive Bayes algorithms. Other packages that use MCMC methods such as *Hydra* (Warnes, 2002) and *Blaise* (Bonawitz *et al.*, 2007) are mostly limited to a single and widely-used MCMC method called the Metropolis-Hastings (MH) sampler, which often performs well in small networks (Su and Borsuk, 2016). However, with larger networks it is slow in mixing and convergence, and often gets trapped in local optima (locally high probability networks) which results in failure to find true BN structures (Su and Borsuk, 2016). Moreover, many of these BN packages require basic programming skills (e.g. Java, C++) (Murphy, 2007).

Learning BN structures based on neighbourhood searches has been used in the literature by different heuristic samplers e.g. Hill-Climbing and Tabu Search (Glover, 1989, 1990). A new version of the Neighbourhood Sampler (NS) developed in (Keith *et al.*, 2008) based on MCMC sampling has not been implemented to infer BN structures until the recent work done by (Alyami, 2017; Alyami *et al.*, 2016a,b). The sampler has been shown to traverse the BN space with reduced chance of getting trapped in local modes. This is partly because the NS possesses a reduction step in which rejected BNs are excluded from being chosen a second time (Alyami *et al.*, 2016b). Each new BN is chosen in two steps: starting from an initial BN *X*, a neighbour *Y* of *X* is first selected, then a neighbour *Z* of *Y* is proposed for final sampling. These two steps help to ameliorate the problem of local modes (Alyami *et al.*, 2016b). Another powerful MCMC method which has the potential to substantially resolve the problem of local modes is called the Hit-and-Run (HAR) sampler (Smith, 1984). The HAR sampler has been recently proposed in (Alyami, 2017) to infer BN structures. The sampler enables transitions from current graphs to distant graphs in a single iteration. It iteratively hits a particular graph in the discrete space and then runs along a random path defined as a sequence of adjacent graphs, in which each graph is immediately adjacent to the graph that precedes it. The performance of the HAR sampler has shown to be efficient to learn BNs compared to a range of famous heuristic samplers (Alyami, 2017). The potential of these two MCMC samplers (NS and HAR) is still unexploited by any BN structure learning package.

In this article, we present new software called BNMCMC (**B**ayesian **N**etwork learning with **M**arkov **C**hain **M**onte **C**arlo) featuring NS, HAR and MH samplers to learn BN structures from data. The BNMCMC software aims to address the above-mentioned problems, and facilitate analysis by practitioners from various disciplines to infer Bayesian Networks. Detailed definitions, notations, algorithms and applications related to the BNMCMC software can be found in (Alyami, 2017; Alyami *et al.*, 2016a,b).

## 2 Approach

To infer BN structures from data, the samplers in BNMCMC (NS, HAR, and MH) explore the *discrete* space of candidate BNs. At each sampling iteration, these samplers iterate through all possible neighbourhoods of a particular BN and sample the best scored one among them (Alyami, 2017). The neighbourhood of a particular BN consists of all networks that can be obtained by modifying a pair of nodes by adding, deleting or reversing an edge. Experimental data is used for learning the conditional probability tables (CPTs) of nodes of a candidate BN and a score is evaluated using those CPTs by applying a probabilistic model which refers to the posterior probability of the network given data. In this version of the software we have used the Dirichlet-Multinomial (DM) model to determine scores of candidate BNs. In this model, the Dirichlet distribution describes the *a priori* knowledge regarding the CPTs and the Multinomial distribution describes the likelihood of observed data. To accelerate the sampling process, each sampler adopts adaptive techniques that maintains a list of neighbouring BNs to the current graph, and dynamically updates this list as each new graph is sampled. Next, it calculates the score of a candidate BN in constant time (*O*(1)) with respect to the number of nodes (Alyami, 2017). The architecture and flowchart of BNMCMC is outlined in Fig. 1A.

**Fig. 1.**
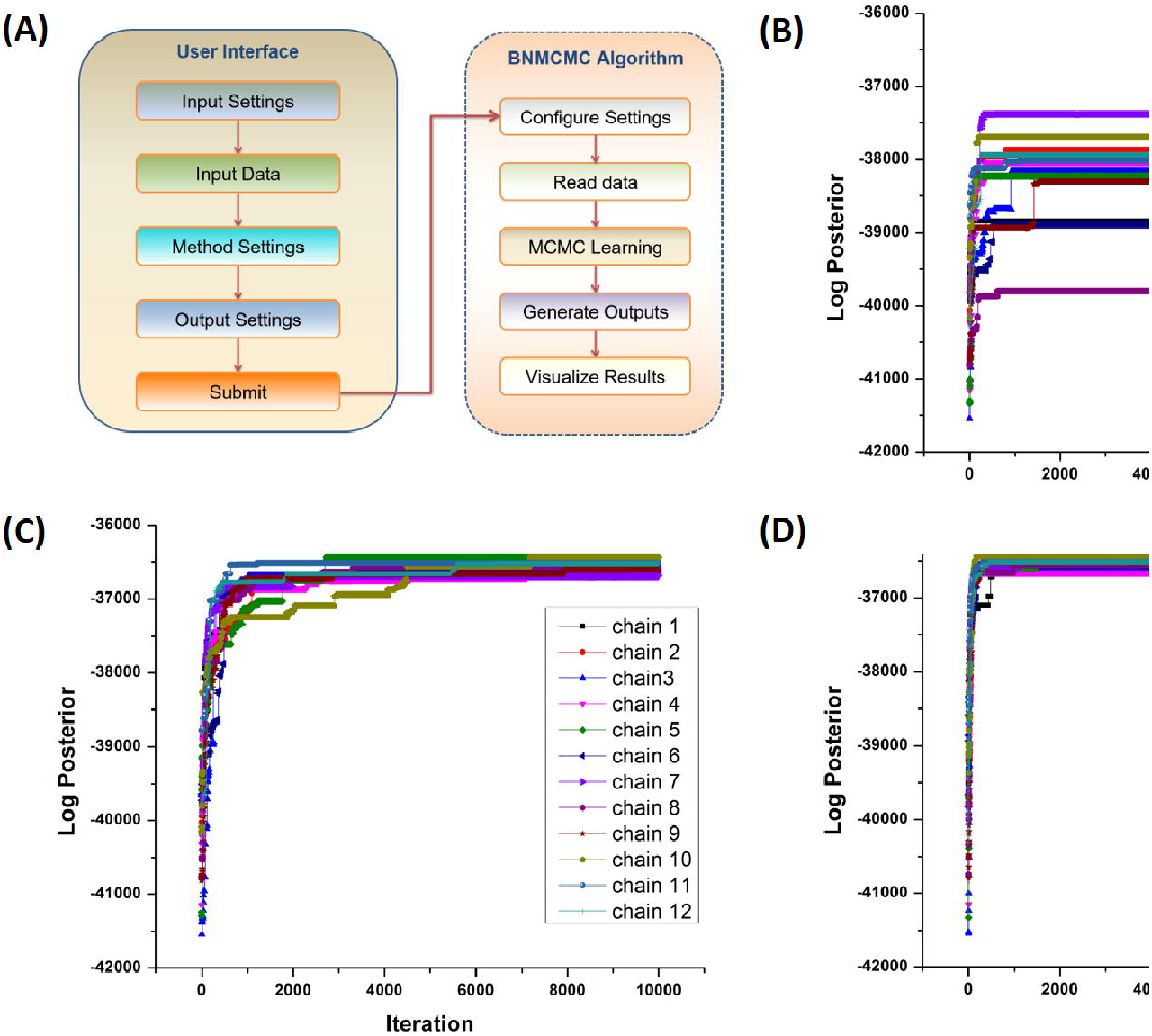
(A) Architecture and flowchart of BNMCMC. Based on the experimental data collected from (Sachs et al., 2005), the log posterior probability of sampled BNs are shown for 12 chains using (B) Metropolis-Hasting (MH), (C) Hit-and-Run (HAR), and (D) Neighbourhood Sampler (NS), where each plot represents a time series indicating sampled networks in multiple runs of each algorithm. It is evident that MH is trapped at local maxima with inferior log posterior probability when compared to HAR and NS.

BNMCMC comes with both standalone and web versions. The BNMCMC web server using ASP.NET under Microsoft Visual Studio platform where C#.net, JQuery and core JavaScript were used for server and client-side coding, respectively. D3.js libary is used for visualizing BNs, which is a very useful tool for zooming in or out large BNs. The standalone version is implemented using C.net. BNMCMC is user-friendly with intuitive software design, requires no programming skills,and provides the following key features:

- It learns BN structures from experimental data (plain.txt format) using three MCMC sampling methods: NS, HAR, and MH, which execute in parallel using multi-threading.
- To increase the responsiveness of BNMCMC server and report the sampling progress, an interactive progress monitor interface is developed with Ajax function calling (from client to server) by following a design pattern proposed by (Mohaimenuzzaman and Rahman, 2015).
- The sampling process can be configured with a set of limited but intuitive parameters, e.g. the number of nodes in BN, number of sampling iterations, the maximum number of parent nodes (Max In-Degree), the maximum number of children nodes (Max Out-Degree), and the length of burn-in interval. Furthermore, HAR sampler requires one special parameter, called *λ*, which indicates the maximum number of edge modifications that may be applied to the current BN.
- It yields two types of results: numerical and graphical. Graphical output includes a scatter plot of log posterior probability of the sampled BN per sampling iteration, and visualizations of two types of inferred BNs: one with edges selected with mean posterior probability *≥* some user-defined threshold, and the other BN that is sampled with highest frequency. Numerical output optionally includes files containing the list of sampled BNs with their corresponding frequencies and their log posterior probabilities (also showed as graphical output).

## 3 Application - Inferring Raf signalling pathway

To demonstrate BNMCMC, we inferred the causal structure of the Raf signalling network from experimental data. Raf signalling structure was first studied in (Sachs *et al.*, 2005), but has been further enhanced over the past two decades. The original network contains 11 nodes, where the nodes are phosphoproteins and the causal edges indicate signal transduction events. For each node, 5400 continuous samples (without missing data) were collected from (Sachs *et al.*, 2005) and the sample gene expression values were discretised into three categories: high, medium and low (Sachs *et al.*, 2005).

Using the above dataset we ran NS, HAR and MH samplers with 10,000 iterations (5000 burn-in), maximum out-degree = 6, maximum in-degree = 3, as in (Sachs *et al.*, 2005). We used a DM distribution function to calculate the posterior probability of a candidate BN. We repeated the above experiment in 12 independent chains (with 12 random initial graphs) for which results are shown in supplementary Table 1. Figure 1B-D show the time series of log posterior probabilities of sampled BNs for multiple chains in MH, HAR and NS samplers, respectively. Note that the MH sampler became trapped at different local modes of the log posterior distribution for each of the 12 Markov chains. This problem was not avoided even with a much larger number of iterations (Alyami, 2017). However, in contrast, the HAR and NS samplers produced higher posterior probabilities and convergence towards a common value for all 12 Markov chains. More detailed results including the inferred BN structures for all samplers and for all 12 chains are provided in supplementary data.

